# Investigating cortical buffering effects of acute moderate and light intensity exercise: a cTBS study targeting the left dorsolateral prefrontal cortex

**DOI:** 10.1101/2020.09.14.297010

**Authors:** Felicia Manocchio, Cassandra J. Lowe, Peter A. Hall

## Abstract

**Background:** The beneficial effects of both single-session bouts of aerobic exercise and therapeutic exercise interventions on the cortical regions associated with executive functions (i.e., prefrontal cortex (PFC)) have been well documented. However, it remains unclear whether aerobic exercise can be used to offset temporary fluctuations in cortical activity.

**Objective:** The current study sought to determine whether a single session of moderate intensity aerobic exercise can offset the attenuating effects of continuous theta burst stimulation (cTBS) targeting the dorsolateral prefrontal cortex (dlPFC).

**Methods:** Twenty-two right-handed participants between 18-30 years completed a 20 minute session of light intensity (10% heart rate reserve (HRR)) and moderate intensity (50% HRR) exercise in a counterbalanced order. Following each exercise session, participants received active cTBS to the left dlPFC. Changes in executive functions were quantified using a flanker paradigm employed at baseline, post-exercise and post-cTBS time points. Additionally, EEG methodologies were used to measure changes in inhibitory control specific event-related potential components (i.e., P3 and N2) in response to the flanker task.

**Results:** Behavioural results from the flanker task revealed a significant improvement in task performance following an acute bout of moderate intensity exercise. Furthermore, the effect of cTBS in both light and moderate intensity conditions were non significant Similarly, EEG data from P3 and N2 ERP components revealed no changes to amplitude across time and condition. P3 latency data revealed a significant effect of time in the light intensity condition, such that latency was faster following cTBS. Similarly, latency data within the N2 ERP component revealed a significant effect of time on congruent trials in the light intensity condition; N2 latency was faster following cTBS.

**Conclusion:** The current study revealed that light and moderate intensity exercise may provide a buffer to cTBS-induced attenuation of the dlPFC. This study provides empirical and theoretical implications on the potential for exercise to promote cognitive control.

## Introduction

Across the lifespan, executive functions are a robust predictor of several psychosocial, intellectual, and health-related outcomes, including academic performance, the adherence to health protective behaviours, and substance use and abuse (Moffitt et al., 2011; Kim et al., 2011). Indeed, the consistent implementation of several everyday behaviours, such as dietary self-regulation, medication adherence, and academic performance (Diamond, 2013; Insel, et al., 2006; Lowe, Staines & Hall, 2019), are thought to be dependent on the higher-order cognitive processes typically associated with executive functions, and by extension the underlying cortical substrates (i.e., the prefrontal cortex). Although executive functions are relatively stable within individuals, several naturally occurring variables can induce state like fluctuations in prefrontal cortical activity, leading to discernable reductions in cognitive control. These modulators include, but are not limited to; sleep restriction, acute stress, and alcohol intoxication (Arnsten, 2009; Lowe, Safati, & Hall, 2017; Marinkovic, Rickenbacher, Azma, & Artsy, 2012; Schulte et al., 2012). The impact of these modulators on executive functioning are of central concern, as such flucuations can increase the likelihood individuals will engage in maladapative behaviours, including overconsuming calorie-dense foods or partaking in risk-seeking behaviours (Brondel, Romer, et al., 2010; Lowe, Hall, & Staines, 2014; Lowe, Staines, et al., 2018; Romer et al., 2009). Over time, this can result in profound implications for individual health and well-being. Considering the crucial role of optimal executive functioning in daily living, and the commonality of executive function disrupters, determining methods to support and maintain optimal cognitive functioning is therefore of the utmost importance

Broadly concieved, the term executive function is used to denote a set of higher-order cognitive processes that play a critical role in “top-down” control and goal-directed behaviours (Baddeley, 1996; Miyake et al., 2000); these processes are distinguishable from other crystalized forms of mental activity. Constituent regions within the executive control network, including the prefrontal cortex (PFC), posterior parietal cortex, inferior frontal junction, and anterior insula comprise the neuroanatomical regions typically associated with executive functioning (Alvarez & Emory, 2006; Derrfuss, Brass, Neumann, & von Cramon, 2005; Hutchison & Morton, 2016; Miller & Cohen, 2001; E. K. Miller, 2000; Niendam et al., 2012; Yuan & Raz, 2014). Within this network, the PFC, particularly the dorsolateral prefrontal cortex (dlPFC), is thought to be the neuroanatomical region most central to executive control. For instance, increased BOLD activation within the dlPFC is observed when individuals complete numerous executive function tasks, including those designed to assess working memory, inhibitory control or response interference, and task-switching (Barbey, Koenigs, & Grafman, 2013; Nee, Wager, & Jonides, 2007). Hemispheric lateralisation within the PFC is thought to contribute to distinct facets of response inhibition and working memory. Specifically, leftwards activation is often associated with tasks involving the top-down control of attentional sets (Vanderhasselt, De Raedt, & Baeken, 2009), and verbal working memory demands (Wager & Smith, 2003), whereas motor response inhibition (Aron, Robbins, & Poldrack, 2014), visual-spatial working memory (Smith, Jonides, Koeppe, 1996; Reuter-Lorenz et al., 2000) and macro-adjustments of cognitive control (Vanderhasselt et al., 2009) appear to be more dependent on rightward activation.

Given the association between dlPFC functionality and executive functions, targeted interventions aimed at enhancing dlPFC functionality can potentially support and maintain optimal executive functioning. Such interventions may be especially pertinent in reducing the impact of acute modulators on executive control. Along these lines, a growing body of evidence has demonstrated that both long-term randomized interventions and acute bouts of aerobic exercise can improve executive function task performance (Chang, Labban, Gapin, & Etnier, 2012; Ludyga, Gerber, Brand, Holsboer-Trachsler, & Pühse, 2016; Smith et al., 2010, Chaddock-Heyman, et al., 2013). These effects are observed following as little as 20 minutes of moderate intensity aerobic exercise, with the largest effects being observed 11-20 minutes post-exercise (Chang et al., 2012), and are apparent for up to 50 minutes post-exercise (Joyce, Graydon, McMorris, & Davranche, 2009).

Although, a general improvement in cognitive functioning is observed following acute bouts of aerobic activity, the largest effects are observed on measures of executive functions (McMorris & Hale, 2012), indicating that executive functions may be the most sensitive to exercise-induced improvements in neurocognitive functioning. Within the acute exercise and executive function literature, the majority of studies have focused predominately on the behavioural inhibition subcomponent. Collectively, these studies have demonstrated that a bout of acute aerobic exercise can improve task performance on measures of inhibitory control in children, young adults and older adults (Barella, Etnier, & Chang, 2010; Chang, Chu, Wang, Song, & Wei, 2015; Hyodo et al., 2016, 2012; Kamijo et al., 2009; Lowe, Hall, Vincent, & Luu, 2014; Lowe, Kolev, & Hall, 2016; Lowe, Staines, & Hall, 2016; Petrella et al., 2018; Samani & Heath, 2018; Yanagisawa et al., 2010). Taken together, the available evidence seems to suggest that acute bouts of aerobic exercise can facilitate improvements in inhibitory control.

The incorporation of neuroimaging methods into exercise interventions have provided important insight into the neural processes underlying the observed exercise-induced enhancements in executive functions. Electrophysiological studies, in particular, have consistently demonstrated that the latency and amplitude of the P3b and N2 event related potential (ERP) components are especially sensitive to aerobic exercise protocols. While the P3b is typically regarded as reflecting the allocation of attention resources, the N2 is more often regarded as a measure of conflict monitoring and response inhibition (Folstein & Van Petten, 2008; Polich, 2007). More specifically, the P3b is an endogenous component generated by an anatomically diffuse network of cortical and subcortical structures. This component is thought to reflect the neural processes underlying several higher cognitive functions typically engaged during executive control tasks (e.g., inhibition, working memory, attention, and stimulus processing; Polich, 2007). The amplitude of the P3b is thought to reflect the neural or attentional resources afforded to a given task or stimulus, whereas, the latency is thought to reflect processing speed (Polich, 2007).

The functionally distinct N2 component is commonly observed in conjunction with the P3b (Folstein & Van Petten, 2008). The N2 is a frontocentral component that is thought to be generated by medial-frontal and latero-frontal neuroanatomical regions (Karch, Feuerecker, Leicht, et al., 2010). The N2 amplitudes are thought to represent the capacity to control incorrect responses and otherwise exert cognitive control during the early stages of response inhibition (Folstein & Van Petten, 2008; Tillman & Wiens, 2011; Veen & Carter, 2002). Thus, the N2 amplitude appears larger (i.e., more negative) when an individual exerts inhibitory control (i.e., for incongruent compared to congruent trials; (Bartholow et al., 2005; Heil, Osman, Wiegelmann, Rolke, & Hennighausen, 2000; KOPP, RIST, & MATTLER, 1996; Tillman & Wiens, 2011; Yeung, Botvinick, & Cohen, 2004).

Across a variety of cognitive tasks, greater amounts of physical activity are associated with larger P3b amplitudes and shorter latencies, indicating that the habitual engagement in physical activity has beneficial effects on neurocognitive functioning that manifests as increased attentional control and faster processing speeds (Hillman, Castelli, & Buck, 2005; Hillman, Belopolsky, Snook, Kramer, & McAuley, 2004; Polich & Lardon, 1997; Pontifex, Hillman, & Polich, 2009). Such physical activity-related modulation of cortical functionality is substantially larger for cognitive measures requiring greater amounts of executive control (Hillman et al., 2006). Similarly, exercise-induced enhancements in P3 amplitude (Chang et al., 2017; Drollette et al., 2014; Hillman et al., 2009; Hillman, Snook, & Jerome, 2003; Kamijo et al., 2009) and reductions in P3 latencies (Chang et al., 2017; Drollette et al., 2014; Hillman et al., 2003; Kamijo et al., 2009) are observed following acute bouts of aerobic exercise, and these effects are specific to those task components involving some degree of inhibitory control. Together, these data suggest that aerobic exercise generally improves neurocognitive functioning by increasing inhibitory and attentional control abilities, and cognitive processing speed during stimulus encoding.

Expanding on this work, Lowe and colleagues (2017) recently demonstrated that aerobic exercise has the capacity to promote cortical resilience (i.e., the ability of brain tissue to rebound from temporary perturbations in a short period of time; Hall, Erickson, Lowe, & Sakib, 2020) in healthy young adults. In this study, inhibitory repetitive transcranial magnetic stimulation (rTMS; specifically, continuous theta burst stimulation (cTBS)) was used attenuate cortical activity in neuronal populations underlying the left dlPFC. The application of cTBS induces temporary fluctuations in cortical excitability for up to 50 minutes following stimulation (Huang, Edwards, Rounis, Bhatia, & Rothwell, 200; Wischnewski & Schutter, 2015). This was followed with either an acute bout of moderate intensity aerobic exercise (50 % heart rate reserve) or light intensity exercise (10% heart rate reserve). Results indicated that dlPFC function recovered at a significantly faster rate in moderate intensity relative to very light intensity exercise condition. Specifically, at the 40-minute post-stimulation point, 101.3% of the attenuation in Stroop performance was recovered in the moderate exercise group, but only 18.3% in the light intensity exercise control group, demonstrating that aerobic exercise can promote cortical resilience. These findings have important theoretical and empirical implications for how we conceptualize the neuroprotective effects of exercise and supports the use of therapeutic approaches to improving cognitive function.

Currently, there is evidence to suggest that a single bout of moderate intensity exercise can promote cortical resilience in the PFC (Lowe, Hall, Staines, 2017) as well as the motor cortex (Singh, Duncan, Staines, 2016). However, the notion of whether exercise can act as a buffer (i.e., to reduce the attenuating effect of cTBS) for regions that support executive control (i.e., dlPFC) has been largely unexplored. As such, the overall purpose of the current study was to test the perturbation-buffering potential of an acute bout of aerobic exercise. It was hypothesized that moderate intensity exercise would reduce cTBS-induced attenuation of dlPFC function, compared to light intensity exercise. It was further hypothesized that such attenuation would be evidenced on behavioral (i.e., task performance) and electrophysiological (i.e., ERP components, P3 and N2) indicators of executive control.

## Methods

A total of 22 healthy young adults, aged 18 to 30 years old (8 women; 21.8 ± 2.89 years old) participated in this study. Participants were recruited from undergraduate courses using an online recruitment system, and from recruitment posters located around the university campus. All participants indicated that they were naïve to TMS, right-handed, and had normal or corrected-to-normal vision. Participants were financially compensated for their participation. Study procedures were approved by the University of Waterloo Research Ethics Committee and conformed to the ethical standards outlined the Declaration of Helsinki. Informed consent was obtained from all participants prior to the start of the study protocol.

Prior to participation, participants were screened to be free of any neurological or psychiatric, and physical health conditions that may increase the risk of an adverse reaction to either the TMS or aerobic exercise protocol. Participants were excluded from the study if they had a) been diagnosed with a neurological or psychiatric condition (i.e., epilepsy, depression, anxiety), b) being treated with any psychiatric medications; c) had a family history of epilepsy or hearing loss; d) history of head trauma (i.e., concussion); e) experienced chronic headaches or migraines; f) has metal in the cranium and/or any implanted electronic or medical devices (i.e., electronic pacemaker, implanted medication pump); g) were pregnant; h) answered “yes” to any of the questions of the Physical Activity Readiness Questionnaire (PAR-Q).

### Procedure

A within-subject study design was utilized, such that participants completed both the moderate intensity and light intensity exercise conditions in a counterbalanced order; exercise sessions were separated by a one-week intersession interval. Study sessions were identical, with only the exercise intensity varying.

Upon arrival, participants were fitted with the EEG cap. Next, they were asked to complete the Profile of Mood States-2 Adult Short (POMS-2A; Heuchert & McNair, 2014), and flanker task. Following baseline assessments, participants completed one session of either light intensity or moderate intensity exercise in a counterbalanced order. Following a 10-minute post-exercise interval, participants completed the POMS-2A and flanker task (post-exercise). Next, cTBS was applied over the left dlPFC, and then participants completed the POMS-2A and flanker task (post-cTBS) again. At the end of the second study session participants were asked to complete a series of questionnaires pertaining to demographics, dietary habits, and physical activity patterns.

### Flanker Paradigm

A modified version of the Eriksen flanker task (Eriksen & Eriksen, 1974) was used to quantify the exercise and cTBS effects on executive functioning. In this version, a fixation cross followed by a five-letter string, composed of Hs and/or Ss, was presented focally on a computer screen. Participants were instructed to indicate whether the center target stimulus among four identical congruent (i.e., HHHHH or SSSSS) or incongruent (i.e., HHSHH or SSHSS) was a “S” or a “H” as quickly and accurately as possible. Responses were made using the associated keys on a keyboard. For each trial, the flanker stimulus was presented for 500ms, followed by a random inter trial interval (ITI) between 600 and 1000ms. Participants completed a total of 240 trials with equiprobable congruency, such that half the trials consisted of congruent stimuli (120 trials) and the other half incongruent stimuli (120 trials). The primary dependent measures were task accuracy, and the flanker interference score (reaction time on correct incongruent trials minus correct congruent trials); higher interference scores are indicative of poorer inhibitory control.

### Aerobic Exercise Protocol

For both the moderate and light intensity exercise conditions, participants were asked to walk on a treadmill for 20 minutes. Speed and incline were gradually adjusted until participants reached their target heart rate (THR), determined using the Karvonen formula and heart rate reserve (HRR; the difference between the age-predicted heart rate max [220-age] and RHR): THR= RHR+ (Target intensity*([220-age]-RHR). Target intensity was set at 50 percent HRR for the moderate intensity condition, and less than 10 percent HRR for the light intensity condition.

### Theta Burst Stimulation Procedure

Continuous TBS was administered using a 75 mm outer diameter figure-8 coil (MCF-B65) connected to a MagPro (model X100) stimulation unit (Medtronic, Minneapolis, MN, USA). Coil positioning was monitored using a computerized frameless stereotaxic system and neuronavigation software (*Brainsight TMS*, Rogue Research, Montreal, Canada) in conjunction with an T1 weighted structural MRI scan, normalized to MNI space, from a previous data set (the same scan was used for all participants). The left dlPFC was located using the International 10-20 system (Herwig, Satrapi, & Schönfeldt-Lecuona, 2003). Consistent with prior work (Bolton & Staines, 2011; Lowe, Hall, & Staines, 2014; Lowe, Staines, et al., 2016, 2018), cTBS - a 40 second continuous train consisting of 600 pulses applied in the theta burst pattern (bursts of three stimuli at 50 HZ repeated at 5 Hz frequency; (Huang et al., 2005) - was administered over the left dlPFC by positioning the coil at a 9O°angle from the mid-sagittal line with its center positioned over F3. Stimulation intensity was set at 80% resting motor threshold (RMT). RMT was defined at the lowest stimulation intensity required to produce a motor-evoked potential (MEP) with a peak-to-peak amplitude exceeding 50 μV in at least 5 out 10 consecutive trials. Stimulation was applied over the contralateral motor cortex, at a 45° angle tangentially to the scalp, with the handle pointing posteriorly. For each study session, individual RMTs were determined using electromyography measured from the right abductor pollicus brevis (APB) muscle.

Average cTBS stimulation intensity (% stimulator output) for the moderate intensity and very light intensity exercise condition were 53.41(*SD*=6.57) and 53.40 (5.63) respectively. No significant differences in cTBS stimulation intensity between exercise conditions was observed (*t*(21)=.012, *p*=.990,95 % CI [-1.58, 1.60]).

### EEG recording and analysis

Continuous EEG data were recorded from 10 midline and frontal electrode sites [FP1, FP2, FPz, Fz, F3, F4, FCz, Cz, CPz, Pz] using a 64 Ag/AgCI electrode Neuroscan Quick-Cap (Compumedics, Charlotte, NC), referenced online to a mid-line electrode located between Cz and CPz and grounded to AFz. Data were sampled and digitized at a rate of 1000 Hz with a bandpass filter of .1 to 70 Hz. All channel recordings had impedance values below 5kΩ, and impedance was monitored before and after cTBS and exercise.

Offline, data were pre-processed using MATLAB 2018a and the EEGLAB toolbox (version 14; (Delorme & Makeig, 2004). EEG data were filtered using a 50 HZ low-pass filter to remove signal drifts and line noise, and re-referenced to the bilateral mastoids (M1, M2). Response-locked ERP segments spanning 100 to 800 ms post-stimulus onset for correct congruent and incongruent trials were computed individually for each participant. The resulting data were individually decomposed using temporal independent component analysis (ICA) with extended infomax algorithm (Bell & Sejnowski, 1995; Delorme, Sejnowski, & Makeig, 2007). Independent components that were not located within the cortex, as well as those components elicited by spurious movement and ocular artifacts were removed. Afterwards, data were visually inspected, and residual nonstereotyped artifacts were removed.

For all dependent measures [incongruent, congruent], data were averaged relative to a 100 ms pre-stimulus baseline. As recommended by Luck and Gaspelin (Luck & Gaspelin, 2017), measurement time windows and electrode regions of interest for each component were determined *a priori* using typical electrode sites and time windows from prior studies. This reduces type 1 error and the bias towards significance. Electrode sites and measurement time windows were confirmed via visual inspection of the grand average waveforms. Data for all components was extracted from midline electrode sites Fz, FCz, CZ, and Pz..

Stimulus-locked amplitude and latency measures for each ERP component was calculated by determining the peak amplitude (μV) for correct congruent and incongruent flanker trials within two-time windows. The P3b was defined as the peak amplitude between 300-600ms post-stimulus presentation. Likewise, the flanker N2 was defined as a negative deflection peaking between 200 and 300 ms. As recent evidence has suggested that averaging ERP amplitudes across several electrodes to increases signal reliability (Huffmeijer, Bakermans-Kranenburg, Alink, & van IJzendoorn, 2014). As such, amplitude and latency data were averaged across electrode regions of interest for each component. Specifically, a P3b cluster was created by averaging the peak amplitude (μV) and latency across central and parietal electrodes (Cz, CPz, Cz). The P3b is typically maximal at central parietal sites (Polich, 2007).The amplitudes and latencies from frontocentral electrode sites (Fz, FCz Cz) were averaged together to create a frontocentral N2 cluster

### Data Analytic Approach

Accuracy was calculated as the proportion of correct responses to congruent and incongruent trials. Prior to analyses, reaction times less than 100 ms and greater than 3 standard deviations from the individual mean reaction time were excluded. Frequentist analyses were conducted using SPSS (version 25; IBM Corp, Armonk, NY). First, paired sample t-tests were conducted to ensure baseline comparability between conditions in Flanker task performance and cTBS stimulation intensity. Behavioural data were analyzed using a 2 × 3 mixed ANOVA with exercise condition [moderate, very-light] and time [baseline, post-exercise, post-cTBS] as the within subject factor, and order of the exercise condition as the between subject factor. Significant interactions were followed up with simple effect one-way ANOVAS at each level of exercise condition. For significant effects, Fisher’s least square differences (LSD) post-hoc tests were performed. Electrophysiological data were analyzed using separate 2[moderate, very-light] x 3 [baseline, post-exercise, post-cTBS] x 2 [incongruent, congruent] repeated measures ANOVAs for N2 amplitude, N2 latency, P3b amplitude, P3b latency. Significant three-way interactions were followed up with a 3[baseline, post-exercise, post-cTBS] x 2 [congruent, incongruent] repeated measures ANOVAs at each level of exercise [moderate, very-light]. Significant interactions were followed up with simple effect ANOVAs. For significant effects, Fisher’s LSD post-hoc tests were performed.

To account for the small sample size, we also employed a Bayesian approach. Bayesian inference is a model comparison approach that is better suited to accommodating small sample sizes than standard frequentist approaches. Bayesian statistics provide information on likelihood and strength of an effect by evaluating how well the data support the alternative over the null hypothesis. As such, this approach is capable of distinguishing between lack of power and/or precision, and the lack of an effect. Specifically, the Bayes factor quantifies how likely the alternative relative to the null model is. For instance, a Bayes factor (BF_10_) of 10 would suggests strong evidence indicating that the alternative model is 10 times more likely to occur than the null. Bayesian analyses were conducted using the default priors for t-test and ANOVA analyses (Rouder, Morey, Speckman, & Province, 2012) and JASP software using the same procedures described above.

## Results

### Baseline Comparability of Condition

No significant differences in baseline interference scores were observed (*t*(21) = .904, *p*= .376, 95% CI [-6.99, 17.74]; BF_10_=.32l), indicating comparable baseline performance between minimum and moderate exercise conditions. Likewise, no significant differences in cTBS stimulation intensities between exercise conditions were observed (*t*(20)=.01, *p*=.990, 95% CI [-1.58, 1.60]; BF_10_=.228

### Behavioural Results

Performance accuracy, reaction times on incongruent and congruent trials, and flanker interference scores as a function exercise condition (light or moderate intensity exercise) and time (prestimulation, post-exercise, post cTBS) are presented in Table 1. Analyses indicated that the order by exercise condition (*F*(1, 20)=2.15 *p*=.158, *d*=.655), order by time (*F*(2,40)=.73, *p*=.489, *d*=.369), and the three-way [order x time x exercise condition] interaction (*F*(1, 20)=.69, *p*=.507, *d*=.381) were not significant. Although the main effect of exercise condition was not significant (*F*(1, 20)=2.73, *p*=.114, *d*=.739), a significant main effect of time (*F*(2, 40)= 5.43, *p*=.008, *d*=1.04) was observed. This was qualified by a significant time [baseline, post-exercise, post-cTBS] by exercise condition [moderate, very-light] interaction (*F*(2, 40)=5.86, *p*=.006, *d*=1.08).

**Table 1.**
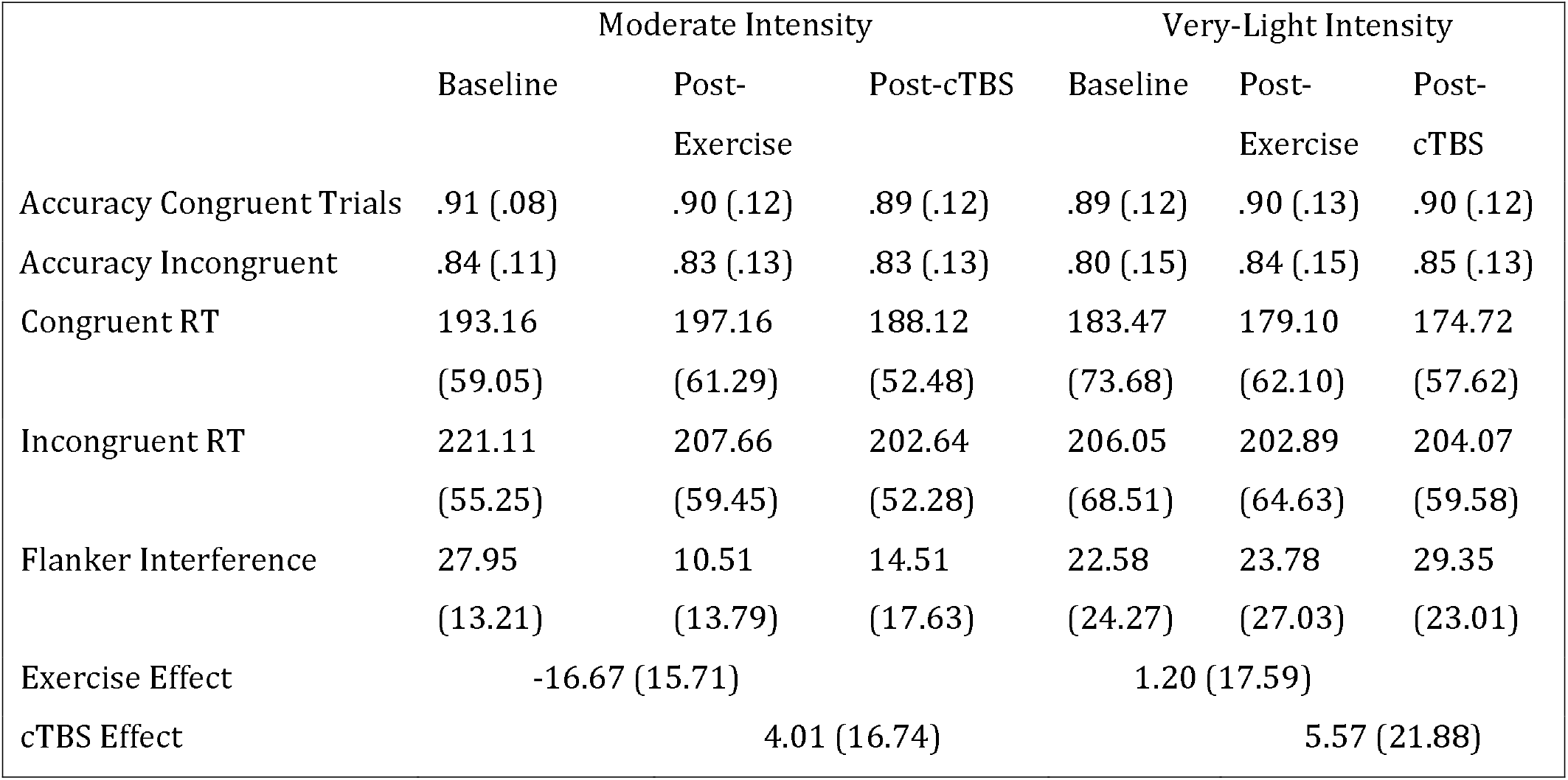
Accuracy, Reaction Time (RT) and Interference Scores Across Time and Condition

Results from the one-way simple main effects ANOVA indicated a significant main effect of time for the moderate intensity exercise condition (*F*(2, 42)=12.23, *p*<.001, *d*=1.526). Planned LSD comparisons revealed that flanker scores were significantly lower (*p*<.001, 95% CI [10.00, 24.88) following the acute bout of moderate aerobic exercise relative to baseline. No significant differences in post-exercise and post-cTBS interference scores were observed (*p*=.274, 95% CI[-11.43, 3.41). Most notably, post-cTBS interference scores were significantly lower than baseline scores (*p*=.003, 95%CI [5.27, 21.60]), indicating a potential buffering effect of aerobic exercise; see Figure 1. The main effect of time was not significant for the very-light intensity exercise condition (*F*(2,42)=1.25, *p*=.298, *d*=.487).

**Figure 1.**
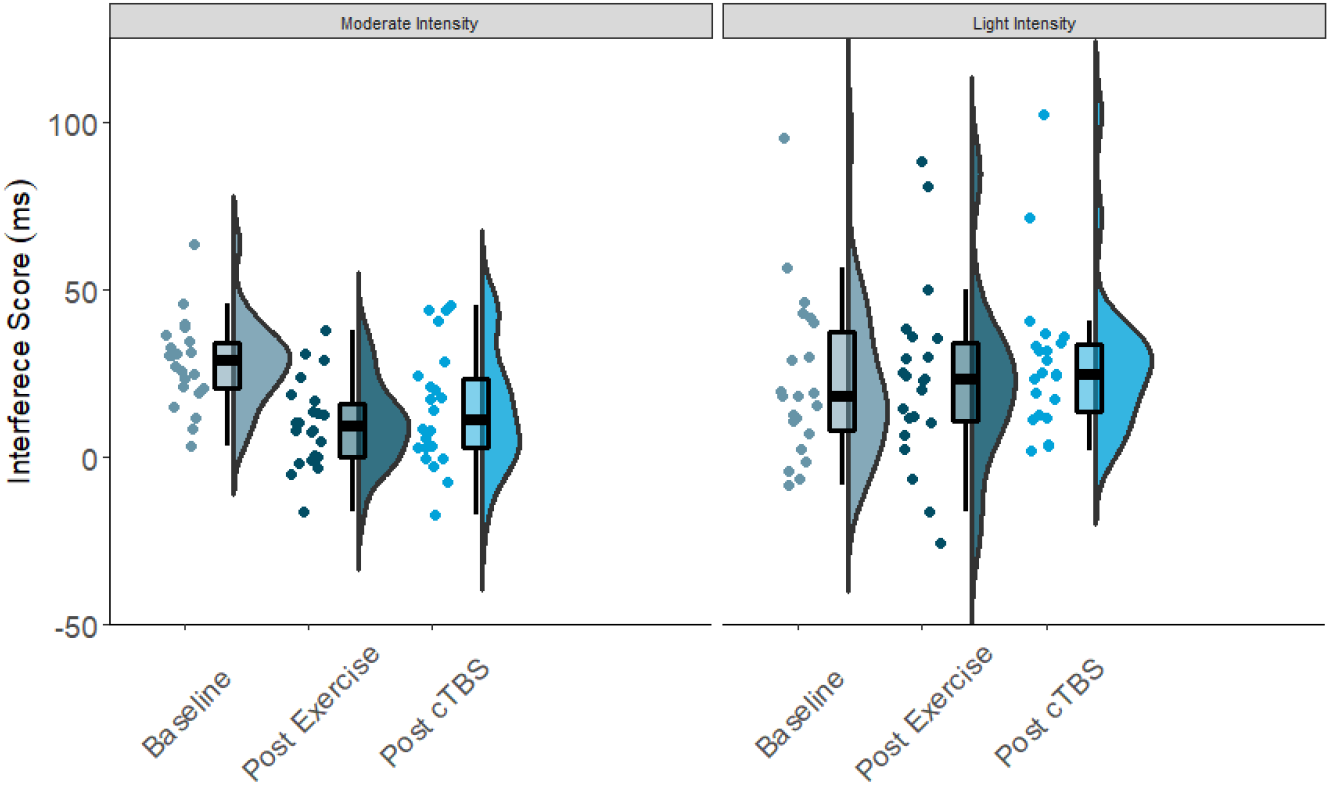
Flanker Interference Scores at Baseline, Post-exercise, Post-cTBS in Light and Moderate Exercise Conditions

**Figure 2.**
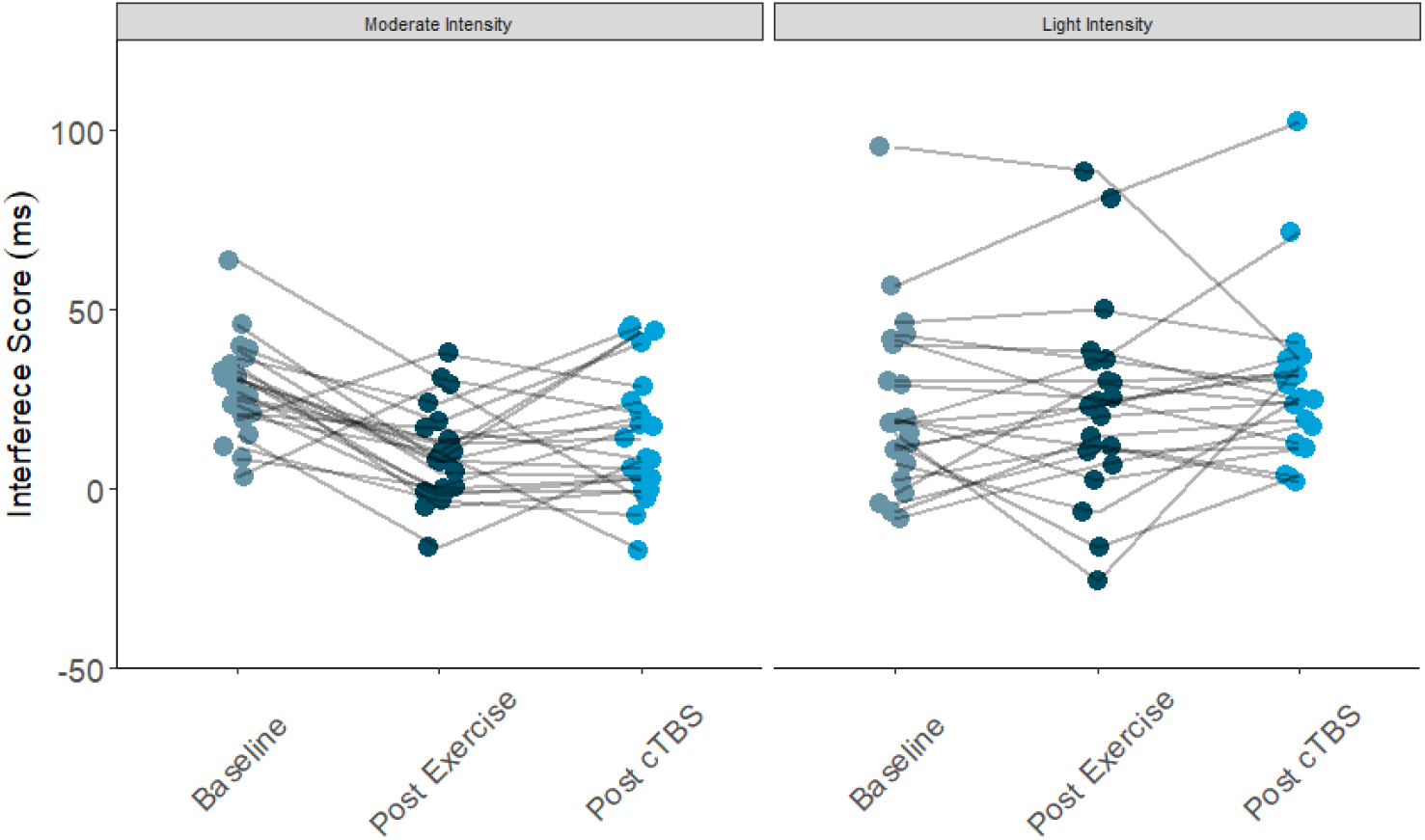
Flanker Interference Scores at Baseline, Post-exercise, Post-cTBS in Light and Moderate Intensity Exercise Conditions

### Bayesian Statistics

Results from the 2 × 3 Bayesian repeated measures ANOVA analyses revealed anecdotal evidence supporting the main effect of exercise (BF_10_= 1.918), and no evidence supporting the main effect of time (BF_10_=.521). Moderate evidence supporting the exercise [moderate, very-light] by time [baseline, post-exercise, post-cTBS] interaction (BF_10_=3.673) was apparent. To assess the evidence supporting the interaction term the Bayes Factor for the model with the interaction term was divided Bayes Factor for the model with only the main effects (exercise and time; 3.673/1.084). The interaction model was preferred the main effects model by a Bayes Factor of 3.39, indicating there was moderate evidence supporting the inclusion of the interaction term over the main effects.

In the moderate exercise condition, the evidence supporting the main effect of time was extremely strong (BF_10_=.936.1180. Post hoc comparisons revealed extremely strong evidence supporting the contention that post-exercise (post-exercise) interference scores were lower than baseline scores (BF_10_=623.512). Likewise, results indicated that there was strong evidence (BF_10_=623.512) supporting the notion that post-cTBS scores differed from baseline. There was no evidence suggesting that post-exercise and post-cTBS interference scores were different (BF_10_=.447). Conversely, there was no evidence supporting a time effect in very-light exercise condition (BF_10_=.261).

## ERP Results

### P3b Amplitude

Analysis of the P3b amplitude revealed a significant main effect of congruency (*F*(1,21)=40.003, *p*<.001, *d*=2.762). Across exercise conditions, the amplitude to incongruent trials was significantly smaller than congruent trials; see Table 2. The main effects of exercise condition (*F*(1,21)=1.85, *p*=.189, *d*=.594) and time (*F*(2,42)=1.04, *p*=.363, *d*=.444), and exercise by congruency interaction (*F*(1,21)=1.32, *p*=.263, *d*=.501) were not significant. Trends towards significance were observed for the exercise by time (*F*(2,42)=2.86, *p*=.069, *d*=.739) and time by congruency (*F*(2, 42)=2.80, *p*=.072, *d*=.732) interactions. The three-way interaction was not significant (*F*(2, 42)=.132, *p*=.876, *d*=.155).

**Table 2.** P3 ERP amplitude and Latencies across condition and time

| **P3b** | **Moderate Intensity** |  |  | **Very-Light Intensity** |  |  |
| --- | --- | --- | --- | --- | --- | --- |
|  | Baseline | Post-exercise | Post-cTBS | Baseline | Post-exercise | Post-cTBS |
| **Amplitude** |  |  |  |  |  |  |
| Incongruent | 7.12 (1.64) | 7.42 (2.30) | 7.13 (1.84) | 5.89 (3.11) | 7.45 (2.57) | 6.44 (3.40) |
| Congruent | 5.56 (1.62) | 5.02 (1.65) | 5.51 (1.27) | 4.41 (1.72) | 5.45 (1.75) | 5.31 (2.23) |
| **Latency** |  |  |  |  |  |  |
| Incongruent | 423.75 (42.86) | 373.78(21.85) | 390.41 (42.01) | 432.10 (47.91) | 412.45(49.04) | 378.62 (41.31) |
| Congruent | 413.30 (419.92) | 401.18 (45.14) | 395.85 (34.30) | 401.12 (50.05) | 413.89 (54.77) | 398.94 (59.13) |

### P3b Latency

Examination into the effects on P3b latency revealed a significant main effect of time (*F*(2, 42) = 8.36, *p*=.001, *d*=1.263). The main effects of exercise condition (*F*(1, 21)=.84, *p*=.370, *d*=.398) and congruency (*F*(1, 21)=.84, *p*=.369, *d*=.398) were not significant. This was qualified by significant time by exercise (*F*(2, 42)=5.28, *p*=.009, *d*=1.003) and time by congruency (*F*(2, 42) =4.96, *p*=.012, n2=.191) interactions. However, the three-way [time x exercise condition x congruency] interaction (*F*(2, 42)=037, *p*=.963, *d*=.090) was not significant.

In the moderate exercise condition, results from the 2 x 3 [congruency by time] ANOVA revealed a significant time by congruency interaction (*F*(2,42)=3.37, *p*=.044, *d*=.800). Similar to the behavioural results, a significant time effect was apparent for incongruent trials (*F*(2, 42)=1 6.720 *p*<.001, *d*=1.784). Specifically, compared to baseline (*M*=423.75, *SE*=9.14) P3b latencies were significantly faster (*p*<.001, 95% CI [33.58, 66.36]) following a bout of moderate intensity exercise (*M*=373.78; *SE*=4.67). Additionally, post-cTBS (*M*=390.41; *SE*=8.96) latencies were marginally higher than those observed post-exercise (post-exercise; *p*=.052, 95%CI [-.18, 33.44). Importantly, post-cTBS latencies were significantly latencies were significantly faster (*p*=.004, 95% CI [12.04, 54.64]) than baseline. No significant time effects were observed for congruent trials (*F*(2,42)=1.004, *p*=.375, *d*=.440).

In the very-light intensity exercise condition, the main effects of time (*F*(2,42)=6.13, *p*=.005, *d*=1.08) and congruency (*F*(2,42) = 8.48, *p*=.008, *d*=1.27) were significant. However, the congruency by time interaction was not significant (*F*(2,42)=1.86 *p*=.168, *d*=.594). Across timepoints, congruent trials had significantly faster latencies than incongruent trials (*p*=.008 95% CI [4.75, 28.50]). In addition, across congruency types, post-cTBS latencies were significantly faster than those observed post-exercise (*p*=.003, 95% CI [9.46, 39.32]) and at baseline (*p*=.006, 95% CI [8.79, 46. 87]). There were no significant differences observed when comparing post-exercise and baseline latencies (*p*=.720, 95% CI [-16.30, 23.19]).

### N2 Amplitude

Results revealed a significant main effect of congruency (*F*(1,21)=13.88, *p*=.001, n2=1.626), such that across exercise conditions and time points the N2 amplitude to congruent trials (*M*=-1.65, *SE*=.09) was smaller than that to incongruent trials (*M*=-2.25; *SE*=.19); see Table 3. The main effects of exercise condition (*F*(1, 21)=.250, *p*=.622, n2=.012) and time (*F*(2,42)=.423, *p*=.658, *d*=.286). No significant interactions were observed (*p*>.20).

**Table 3.**
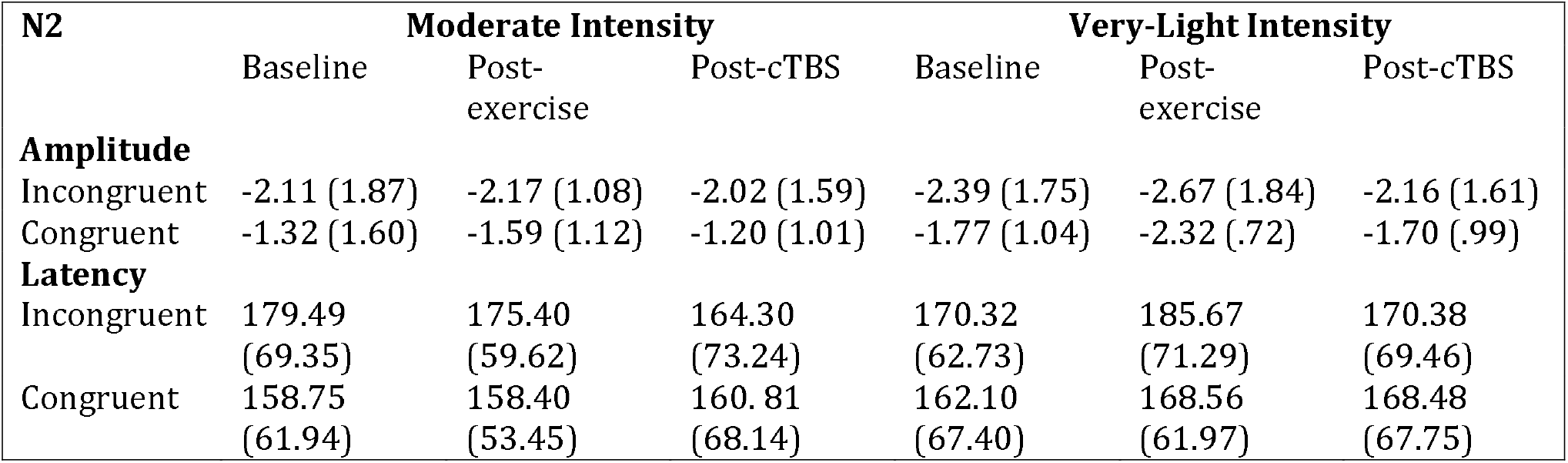
N2 ERP Amplitudes and Latencies Across Condition and Time

### N2 Latency

No significant main effects (*p*>.10) or interactions (*p*>.20) were observed when examining N2 latencies.

## Discussion

The current study sought to assess the capacity of aerobic exercise to offset (i.e., act as a buffer against) the temporary attenuating effects of cTBS targeting the left dlPFC. Overall, findings suggest that both moderate and light intensity exercise provide a protective buffer against temporary perturbations in dlPFC functionality, and by extension executive control. Behaviourally, significant improvements in Flanker task performance were observed following the bout of moderate intensity exercise. Most notably, post-cTBS interference scores did not differ from post-exercise (postexercise) scores, but instead remained significantly lower compared to baseline. Interestingly, there were no significant differences in post-exercise and post-cTBS interference scores in the light intensity exercise condition. While this effect was unexpected, it may suggest that both light and moderate intensity exercise may offset the temporary cTBS induced attenuation to cognitive control, assuming that cTBS was effective in both conditions. Examination of neuroelectric measures revealed shorter P3b latencies to incongruent trials following moderate intensity exercise, suggesting faster cognitive processing speed post-exercise. Similar to the behavioural data, there were no significant differences in post-exercise and post-cTBS latencies. This effect was also apparent in the very-light intensity exercise condition.

Taken together, the current findings add to the well-documented effects of aerobic exercise on cognitive control. Novel to the present investigation is that despite the lack of a discernable exercise effect in the very-light intensity exercise condition, both moderate and very-light intensity exercise seem to provide a protective buffer against experimentally-induced perturbations in cognitive control. This expands on the findings by Lowe, Staines & Hall (2017(Lowe, Staines, et al., 2016)), and provides important insight into the potential of aerobic exercise to act as a protective and recovery mechanism against fluctuations in dlPFC activity. These findings have substantial implications for how we conceptualize the beneficial effects of aerobic exercise on brain health and executive control. That is, while the current study employed an experimental perturbation (specifically, cTBS), there are many naturally occurring perturbations in everyday life, including lack of sleep, mood fluctuations, and acute stress (Porcelli & Delgado, 2009; Shields et al, 2016; Fossati, et al, 2002; Nillson, et al, 2005; Tucker, et al, 2010). If generalizable to these types of everyday perturbations, the current findings suggest the possibility that exercise may serve to provide protection against momentary fluctuations in cognitive control in everyday living. Such findings may have implications for clinical intervention strategies as well. For instance, acute bouts of exercise may provide an optimal intervention or preventative strategy for people who are chronically subject to exposure to lifestyle perturbations mentioned previously (e.g., shift workers, hospital employees). Additionally, given that buffering effects appear to manifest in relation to both moderate and light intensity exercise, the effects of this intervention could extend to individuals who may not be physically able to perform moderate intensity exercise, such as older adults.

While significant behavioural effects were apparent, this did not manifest as changes in P3b or N2 amplitudes, as expected. However, significant P3b latency effects to incongruent trials were observed. Consonant with the behavioural data, shorter latencies were apparent following the bout of moderate intensity exercise. Further, post-cTBS P3b latencies were significantly faster than baseline in the moderate exercise condition. In general, the amplitude of the P3 and N2 ERP components are thought to reflect the neural resources afforded to a given task, whereas, the latency is thought to be an index of cognitive processing speed or the speed at which an individual is able to classify and evaluate a stimulus (Folstein & Van Petten, 2008; Polich, 2007). Therefore, results from the current study seem to suggest that the protective effects of exercise are specific to cognitive processing speeds rather than attentional allocation or conflict adaptation *per say*, a finding that is consistent with several other studies. For instance, P3 latencies in both young and older adults have been shown to be shorter following both light and moderate intensity exercise (Kamijo, 2009, Kamijo, 2007).

Additional evidence supporting this notion comes from rTMS prior studies utilizing neuroelectric measures of cognitive control. Stimulation-induced reductions in the latency of the P3 component is observed following both 10Hz and 20Hz (i.e., excitatory stimulation) rTMS targeting the left PFC (Evers, Böckermann, & Nyhuis, 2001; Pinto et al., 2018; Rektor, Baláž, & Boĉková, 2010). Likewise, several studies using inhibitory rTMS or cTBS targeting the dlPFC have reported that attenuation of left dlPFC activity results in the subsequent increase in P3 latencies (Evers et al., 2001; Hansenne, Laloyaux, Mardaga, & Ansseau, 2004; Jing, Takigawa, Okamura, Doi, & Fukuzako, 2001; Pinto et al., 2018). Further, recent findings from our laboratory demonstrated that stimulation-induced increases in P3b latencies to incongruent Flanker trials are observed following cTBS over the left dlPFC (Lowe, Manocchio, Safati, & Hall, 2018). These findings are noteworthy, as the same stimulation, flanker, and EEG protocols were employed as in the current study. Collectively, these data demonstrate that P3 latencies are sensitive to rTMS-induced attenuation of dlPFC activity, indicating that both moderate and very-light intensity exercises did indeed provide a protective buffer against cTBS-induced perturbations in executive control.

Exercise-induced improvements in inhibitory control are thought to be mediated, at least in part, by increased cerebral blood flow to the left dlPFC (Byun et al., 2014; Endo et al., 2013; Giles et al., 2014; Guiney, Lucas, Cotter, & Machado, 2015; Yanagisawa et al., 2010). There is increasing evidence to suggest that light intensity exercise may also result in increased cerebral blood oxygenation (Byun, et al, 2014). For instance, previous studies have indicated that light intensity walking does indeed increase cerebral blood oxygenation to the PFC (Holtzer, et al, 2011; Suzuki, et al, 2004). Considering the association between aerobic exercise and increased cerebral blood oxygenation, it is possible that both very-light and moderate intensity exercise is sufficient to offset cTBS-induced attenuation neuronal populations within the left dlPFC. It is also possible that elevated levels of certain neurotransmitters such as norepinephrine, epinephrine, serotonin, and dopamine, induced a buffering effect, as they have been demonstrated to be elevated following acute bouts of exercise. ERP latencies and amplitudes are thought to be influenced by dopaminergic function, subsequently impacting cognitive processing speeds and neural resources allocated to a specific task (Ullsperger, et al, 2014). However, the buffering effects of exercise have been largely unexplored and may differ from other proposed mechanisms. Further research is warranted to better determine the neurophysiological processes underlying cortical buffering.

Strengths in the present study include the use of a within-subject study design in an effort to minimize any inter-individual variability. Additionally, the use of cTBS as our neuromodulation protocol provided a safe and reliable method by which to investigate the buffering effects of exercise. Limitations of the present study include sampling a healthy university student population, who may not be as receptive to the effects of exercise compared to an older adult sample due to high initial levels of cognitive performance. Previous literature has suggested that older adults tend to demonstrate greater performance improvements compared to young healthy adults following acute bouts of exercise (Chang, et al, 2012). Finally, the lack of a non-exercise control group served as a limitation. Although both light and moderate intensity exercise appeared to demonstrate a potential buffering effect, this cannot be known definitively in the absence of a no-movement control condition. Without this, an alternative interpretation of the findings is that no cTBS effect emerged in either condition, which is possible given that not all studies show significant perturbation effects on executive function following cTBS. Future studies should aim to disentangle this issue.

## Conclusion

The current study was, to the best of our knowledge, the first to demonstrate the potential protective effects of acute exercise. Findings suggested that acute bouts of both light and moderate intensity exercise may provide a buffer to impairments in cognitive control. Specifically, no significant decrements to performance on the Flanker task were apparent through both behavioural and EEG measures, demonstrated through P3 and N2 ERP components. Findings from this study are noteworthy as it provides theoretical and experimental implications for the therapeutic potential of acute exercise to maintain and support optimal EF. As such, further research examining the buffering effects of exercise is warranted to build on the current findings. Investigating the capacity of exercise to offset attenuation could be beneficial in alternate samples and target groups (e.g., older adults, shift workers). Additional research on the buffering effect of exercise using natural modulators previously mentioned (sleep deprivation, alcohol consumption, acute stress) could provide a deeper understanding into how exercise could impact everyday stressors. Finally, subsequent studies using a no-movement control group could reveal important information regarding the role light intensity exercise plays in reducing attenuation to key areas involved in executive control.

